# Cross-platform comparison of highly-sensitive immunoassays for inflammatory markers in a COVID-19 cohort^1^

**DOI:** 10.1101/2023.10.24.563866

**Authors:** Koji Abe, Joanne C. Beer, Tran Nguyen, Ishara S. Ariyapala, Tyson H. Holmes, Wei Feng, Bingqing Zhang, Dwight Kuo, Yuling Luo, Xiao-Jun Ma, Holden T. Maecker

**Author notes:** Both authors contributed equally to this work (Koji Abe, Joanne C. Beer). Corresponding author: Koji Abe.

## Abstract

A variety of commercial platforms are available for the simultaneous detection of multiple cytokines and associated proteins, often employing antibody pairs to capture and detect target proteins. In this study, we comprehensively evaluated the performance of three distinct platforms: the fluorescent bead-based Luminex assay, the proximity extension-based Olink assay, and a novel proximity ligation assay platform known as Alamar NULISAseq. These assessments were conducted on serum samples from the NIH IMPACC study, with a focus on three essential performance metrics: detectability, correlation, and differential expression. Our results reveal several key findings. Firstly, the Alamar platform demonstrated the highest overall detectability, followed by Olink and then Luminex. Secondly, the correlation of protein measurements between the Alamar and Olink platforms tended to be stronger than the correlation of either of these platforms with Luminex. Thirdly, we observed that detectability differences across the platforms often translated to differences in differential expression findings, although high detectability did not guarantee the ability to identify meaningful biological differences. Our study provides valuable insights into the comparative performance of these assays, enhancing our understanding of their strengths and limitations when assessing complex biological samples, as exemplified by the sera from this COVID-19 cohort.

## Introduction

Multiple commercial platforms offer multiplexed detection of cytokines and related proteins, many using pairs of antibodies to capture and detect target proteins. In Luminex assays, the capture antibodies are coated on microbeads, and detector antibodies are labelled with biotin and then streptavidin-phycoerythrin to yield a fluorescent readout (1). In proximity extension and proximity ligation assays (PEA and PLA) (2), pairs of DNA-tagged antibodies bind to the same target protein molecule and facilitate a ligation or polymerase extension reaction whose product can be detected by polymerase chain reaction (PCR) or sequencing. Previous studies have examined the performance of PEA, known as the Olink assay, in comparison to other platforms, including aptamer-based SomaScan (3–6). These comparisons have demonstrated a range of correlations and highlighted various factors that need to be considered when selecting the appropriate platform. Luminex and Olink assays have also been compared (7), with best correlations observed for the more abundant analytes.

The NIH IMPACC study (IMmunoPhenotyping of A COVID-19 Cohort) evaluated longitudinal clinical and immunological features of 1164 patients hospitalized for COVID-19 from 2020-2022. The study employed multiple highly multiplexed assays for viral, cellular, protein, and genomic phenotyping (8). Among these was the 92-plex inflammation panel for serum cytokines from Olink Proteomics. We sought to test performance characteristics of this assay against Luminex and a novel PLA platform, Alamar NULISAseq (9). Since the latter comprises over 200 targets, it could provide a broader evaluation of cytokines in the progression of COVID-19 in the IMPACC cohort. Our Luminex comparison assay was a set of three panels totalling 80 target proteins, using kits from Millipore Corporation.

For the current evaluation, we chose a small subset of 78 serum samples from 23 IMPACC patients, along with 8 healthy control sera. We performed parallel assays on the Olink, Luminex and Alamar panels, compared the detectability and assessed correlation for common analytes, and compared ability to discriminate COVID-19 from healthy samples, ability to detect changes over time in the COVID-19 patients, and ability to distinguish COVID-19 trajectory groups.

## Materials and Methods

### Cohort

The IMPACC study was conducted to investigate the clinical and immunological characteristics of COVID-19 in hospitalised patients (8, 10). This observational cohort study involved a network of expert clinicians, geneticists, and immunologists with the objective of examining the relationship between the clinical course of COVID-19 and the immune response in racially, ethnically, and geographically diverse adult patient populations across the United States. Serum samples were collected at multiple time points during hospitalization, including enrollment, days 4, 7, 14, 21, and 28. Throughout the hospitalization period, clinical outcome data such as mortality and level of care were diligently documented. For a supplementary analysis, we utilized a framework comprising five distinct disease course trajectories to comprehensively capture the evolution of the clinical course of the disease under investigation. Specifically, trajectory 1 represented a brief length of stay (median 3 days), trajectory 2 denoted an intermediate length of stay (median 7 days), trajectory 3 indicated an intermediate length of stay (median 7 days) with discharge limitations, trajectory 4 corresponded to prolonged hospitalization (median 20 days), and trajectory 5 represented a fatal outcome (8, 11).

### Sample preparation

The IMPACC sample subset used in this analysis included 23 COVID-19 patients with samples collected at up to 6 time points each, giving a total of 78 serum samples (mean age 59.3 years, 6 female; see Table I). An additional 8 serum samples were obtained from 8 healthy adult donors (3 female). All samples were stored at −80°C prior to thawing for each assay. These samples were employed without any filtration or modification. This study was approved by the Stanford IRB.

### Protein analysis methods

Luminex assay (EMD Millipore, Burlington, MA) and Olink immunoassay (Olink Bioscience, Uppsala, Sweden) were performed by the Human Immune Monitoring Center at Stanford University Immunoassay Team. For Luminex assay, Human 80 Plex kits were purchased from EMD Millipore Corporation, Burlington, MA., and run according to the manufacturer’s recommendations with modifications described as follows: H80 kits include 3 panels: Panel 1 is Milliplex HCYTA-60K-PX48. Panel 2 is Milliplex HCP2MAG-62K-PX23. Panel 3 includes the Milliplex HSP1MAG-63K-06 and HADCYMAG-61K-03 (Resistin, Leptin and HGF) to generate a 9 plex. PBS was added to wells for reading in the Luminex FlexMap3D Instrument with a lower bound of 50 beads per sample per cytokine. Custom Assay Chex control beads were purchased and added to all wells (Radix BioSolutions, Georgetown, Texas).

Assay Chex beads were used for non-specific binding correction in the Luminex assay. The procedure encompassed log-transforming the MFI values of both target proteins and CHEX4 beads, followed by mean-centering the log-transformed CHEX4 MFI. Subsequently, a linear regression was performed, regressing protein MFI values against centered CHEX4 MFI values. The resulting non-specific binding-corrected values were derived by adding the regression residuals to the intercept estimate.

For Olink immunoassay, the samples were subjected to Olink inflammatory panel multiplex assay (Target 96 Inflammation), according to the manufacturer’s instructions. The inflammatory panel includes 92 proteins associated with human inflammatory conditions. Briefly, an incubation master mix containing pairs of oligonucleotide-labeled antibodies to each protein was added to the samples and incubated for 16 hours at 4 °C. Each protein was targeted with two different epitope-specific antibodies increasing the specificity of the assay. Presence of the target protein in the sample brought the partner probes in close proximity, allowing the formation of a double strand oligonucleotide polymerase chain reaction (PCR) target. On the following day, the extension master mix in the sample initiated the specific target sequences to be detected and generated amplicons using PCR in 96 well plate. For the detection of the specific protein, Dynamic Array Integrated Fluidic Circuit (IFC) 96×96 chip was primed, loaded with 92 protein specific primers and mixed with sample amplicons including three inter-plate controls (IPS) and three negative controls (NC). Real time microfluidic qPCR was performed in Biomark (Fluidigm, San Francisco, CA) for the target protein quantification. Data were analyzed using real time PCR analysis software via ΔΔCt method and Normalized Protein Expression (NPX) manager. One unit NPX difference amounts to the doubling of the protein concentration.

Alamar’s NULISAseq 200-plex Inflammation panel targets mostly inflammation and immune response-related cytokines and chemokines (9). First, serum samples were centrifuged at 10,000g for 10 min. Supernatant from the serum samples were then analyzed using Alamar’s NULISAseq proteomic platform. For NULISA, the capture antibody is conjugated with partially double-stranded DNA containing a poly-A tail and a target-specific barcode, whereas the detection antibody is conjugated with another partially double-stranded DNA containing a biotin group and a matching target-specific barcode. When both antibodies are incubated with a sample containing the target molecule, an immunocomplex is formed. The formed immunocomplexes are captured by added paramagnetic oligo-dT beads and subsequent dT-polyA hybridization, and the sample matrix and unbound detection antibodies are removed by washing. As dT-polyA binding is sensitive to salt concentration, the formed immunocomplexes are then released into a low-salt buffer. After removing the dT beads, a second set of paramagnetic beads coated with streptavidin is introduced to capture the immunocomplexes in the solid phase a second time. Subsequent washes remove free unbound capture antibodies, resulting in essentially pure immunocomplexes on the beads. Then, a ligation mix containing T4 DNA ligase and a specific DNA ligator sequence is added, allowing ligation of the proximal ends of DNA attached to the paired antibodies and thus generating a new DNA reporter molecule containing unique target-specific barcodes. The levels of the DNA reporter are then quantified by Next Generation Sequencing. Data normalization was done using an internal control spiked into each sample well to remove potential technical variation introduced during data collection. Then data was rescaled and log2-transformed to yield NULISA Protein Quantification (NPQ) units for downstream statistical analyses.

Each sample was measured singlicate in each assay platform.

### Statistical analysis

Data analysis utilized log2 normalized signal intensity across platforms. All observations, including those falling below limit of detection (LOD) or outside of the dynamic range for each assay, were included in the subsequent statistical analyses. The number of common protein targets across platforms were as follows: 33 (across three platforms), 56 (Alamar NULISAseq and Olink Target 96), 58 (Alamar NULISAseq and Luminex H80), and 38 (Olink Target 96 and Luminex H80) (Supplemental Data A). All statistical analyses were conducted using R (version 4.3.0) (12).

### Detectability

To determine detectability, LODs recommended by the respective manufacturers were employed. For all three platforms, LOD is defined as the mean plus 3 times the standard deviation of the blank samples. Detectability was defined as the percentage of measured values out of the 78 total samples above the LOD for each protein target. Since blank samples were not included in the Luminex assay run, LODs were obtained for an independent Luminex run using the same reagent lots.

### Correlation

Since association between platforms are not always linear, nonparameteric Spearman and Kendall correlations were used to assess correlations across platforms using baseline (first-visit) COVID-19 samples. Additionally, to accommodate repeated measures in the entire longitudinal dataset, the R package ’rmcorr’ (13, 14) was utilized to investigate the common within-participant correlation. The rmcorr method is based on a form of ANCOVA and assumes linearity and errors that are independent, identically and normally distributed.

### Differential expression: COVID-19 patients at enrollment versus healthy controls

Differential expression for COVID-19 patients at enrollment (n=23) versus healthy controls (n=8) was assessed using a linear regression model with disease status as predictor of target expression. For each inter-platform comparison, Benjamini-Hochberg false discovery rate (FDR) correction (15) was applied to the relevant subset of targets and significance was set at 5% FDR. Pearson correlation of the estimated coefficients (i.e., log2(fold change)) was calculated for shared targets for each platform pair.

### Differential expression: Change over time in COVID-19 patients

Change in target expression over time in the COVID-19 group (n=23) was assessed using linear mixed effects models with categorical time fixed effects accounting for the 6 time points and a random subject intercept. The joint significance of the time fixed effect was tested using a likelihood ratio test (LRT), as implemented in R package ‘lmerTest’ (16). For each inter-platform comparison, FDR correction was applied to the relevant subset of targets and significance was set at 5% FDR. Pearson correlation of the LRT chi-squared statistics was calculated for shared targets for each platform pair.

### Logistic regression: Predicting COVID-19 outcome using protein expression levels at enrollment

An exploratory analysis was carried out to assess whether protein expression levels at enrollment could predict the most severe (trajectory groups 4 and 5; n=14) versus the least severe (trajectory groups 1, 2, and 3; n=9) COVID-19 patient outcomes. Logistic regression models were fit for each target in each panel using target expression as predictor of the binary trajectory group outcome. Results were evaluated using unadjusted and FDR-adjusted p-values at a 5% significance level.

## Results

### Overall detectability

Overall across-target mean detectability (i.e. percent of signal values above LOD) for the platforms was 94.7% for Alamar NULISAseq (203 targets), 86.5% for Olink Target 96 (92 targets), and 84.1% for Luminex H80 (80 targets).

### Detectability with 33 common targets in 3 assays

The detectability of protein targets varied considerably across the three assays for the 33 shared targets (Figures 1A, 1C). In the case of Alamar NULISAseq and Luminex H80, only 1 of 33 protein targets (CCL3 and IL20, respectively) exhibited ≤ 50% detectability. Olink Target 96 had 7 targets with ≤ 50% detectability (IL13, IL20, IL33, IL4, IL5, LIF, TSLP), with five proteins (IL13, IL20, IL4, IL5, and TSLP) consistently falling below the LOD in at least 75% of the samples. For these 33 targets, mean detectabilities were 96.1% for Alamar, 83.5% for Olink, and 87.2% for Luminex.

**Figure 1.**
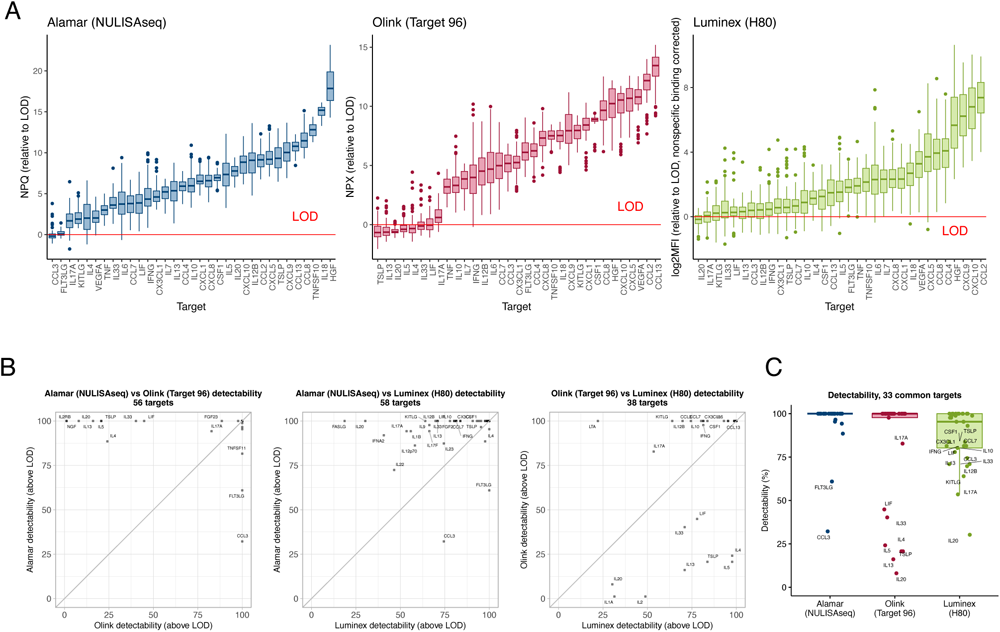
Detectability across platforms. (A) Measured values relative to limit of detection (LOD) for 33 common targets assessed in 78 COVID-19 serum samples from 23 patients and 8 healthy control serum samples. Targets are sorted from left to right based on the median. (B) Comparison of detectability for common targets in each platform pair. (C) Detectability of three platforms for 33 common targets.

### Detectability in pairwise cross-platform comparisons for common targets

Alamar NULISAseq and Olink Target 96 (Figure 1B, left): Of 56 common targets, Alamar showed one target with ≤ 50% of samples above LOD (CCL3), while Olink showed 9 (IL13, IL20, IL2RB, IL33, IL4, IL5, LIF, NGF, TSLP). Alamar mean detectability was 97.3%; Olink mean was 86.5%.

Alamar NULISAseq and Luminex H80 (Figure 1B, middle left): Of 58 common targets, Alamar NULISAseq showed one target with ≤ 50% of samples above LOD (CCL3), while Luminex showed 4 (FASLG, IFNA2, IL20, IL22). Alamar mean detectability was 96.4%; Luminex mean was 85.6%.

Olink Target 96 and Luminex H80 (Figure 1B, middle right): Of 38 common targets, Olink showed 9 targets with ≤ 50% of samples above LOD (IL13, IL1A, IL2, IL20, IL33, IL4, IL5, LIF, TSLP), while Luminex showed 4 (IL1A, IL2, IL20, LTA). Olink mean detectability was 80.5%; Luminex mean was 83.7%.

In summary, in pairwise cross-platform detectability comparisons for common targets, Alamar showed the highest detectability, followed by Luminex and then Olink. The non-specific binding correction improved detectability for Luminex (Supplemental Figures 1A, 1B). Without the correction, Luminex detectability was below Olink in all comparisons.

### Correlation with 33 common targets in 3 assays

We compared Kendall’s coefficient of concordance for 33 common targets using baseline COVID samples (Figure 2A). Alamar and Olink showed moderate to high level of concordance (Kendall’s coefficient mean: 0.56, median: 0.71, range: -0.14 to 0.91). Alamar and Luminex showed a moderate to low level of concordance (Kendall’s coefficient mean: 0.48, median: 0.41, range: -0.30 to 0.85), which was lower compared to Alamar and Olink. Olink and Luminex showed a low level of concordance (Kendall’s coefficient mean: 0.36, median: 0.42, range: -0.16 to 0.82), which was lower compared to both Alamar and Olink, and Alamar and Luminex. Overall, the results suggested varying levels of concordance between different measurement methods. Alamar and Olink exhibited the highest concordance, followed by Alamar and Luminex, and then Olink and Luminex.

**Figure 2.**
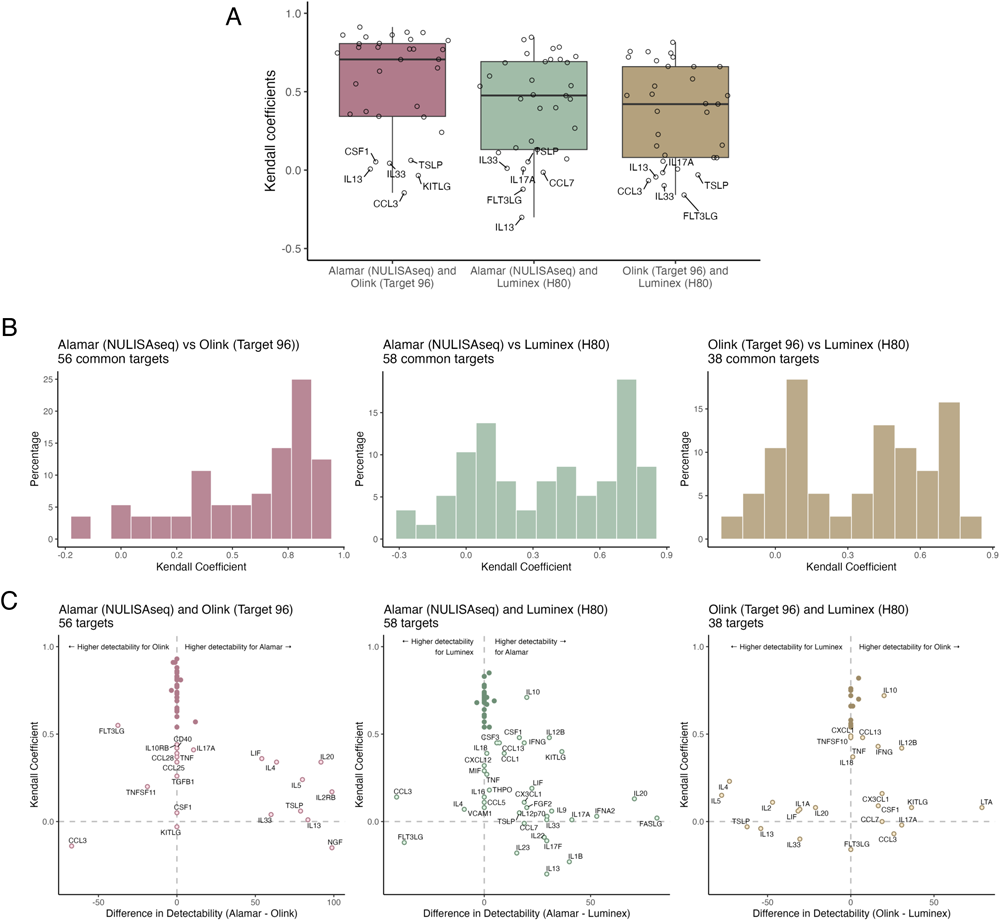
Correlation of measured values across platforms using baseline (Visit 1) samples for COVID-19 (n=23) and Healthy Controls (n=8). (A) Kendall coefficients of each platform pair for 33 common targets. Alamar-Olink showed the highest median correlation, Alamar-Luminex showed second-highest, and Olink-Luminex showed the lowest. (B) Kendall coefficient distributions for all common targets in each platform pair. (C) Association between the difference in detectability (x-axis) and Kendall correlation (y-axis) for common targets in each platform pair using baseline samples. Targets falling on the vertical dashed line showed zero difference in detectability; horizontal dashed line indicates correlation coefficient of zero. Labels highlight protein targets with Kendall coefficient below 0.50 or a difference in detectability greater than 15%. For the most part, larger differences in detectability between platforms corresponds to lower correlation. However, some targets with no detectability difference still showed poor correlation, for example, CSF1 and KITLG for Alamar-Olink, and FLT3LG for Olink-Luminex.

### Correlation in pairwise cross-platform comparisons for common targets

Figure 2B displays the distribution of Kendall’s coefficient in each platform comparison for all of the shared targets. Specifically, the comparison between Alamar and Olink (56 targets) indicated a moderate to high level of concordance (Kendall’s coefficient mean: 0.56, median: 0.70, range: -0.15 to 0.93; Figure 2B, left). The Alamar and Luminex comparison (58 targets) suggested a moderate to low level of concordance (Kendall’s coefficient mean: 0.36, median: 0.40, range: -0.30 to 0.85; Figure 2B, middle), which was lower compared to Alamar and Olink. Moreover, findings indicated a similar level of concordance between Olink and Luminex (38 targets; Kendall’s coefficient mean: 0.35, median: 0.40, range: -0.16 to 0.82; Figure 2B, right) when compared to Alamar and Luminex. Taken together, these results highlighted varying levels of concordance between different measurement methods, mirroring the observation of the highest concordance by Alamar and Olink in comparison involving 33 shared targets.

### Association between detectability and correlation

The association between the difference in detectability and the correlation is depicted in Figure 2C. Notably, in each cross-platform comparison, strong correlations were observed when the differences in detectability were minimal. Conversely, larger disparities in detectability generally corresponded to lower correlation. Furthermore, Alamar NULISAseq exhibited greater detectability than Olink Target 96 and Luminex H80 for targets exhibiting lower overall correlation (more targets to the right of the dotted vertical line at zero in Figures 2C left and middle).

### Differential expression in COVID-19 patients versus healthy controls, 33 common targets in 3 assays

For the 33 common targets, 6 were differentially expressed in all 3 panels (Figure 3A and 3B). All of these targets were upregulated (CCL8, CXCL10, HGF, IL10, IL18, IL6). Alamar uniquely identified upregulated CX3CL1 and LIF, and downregulated IL4. Olink found 9 unique differentially expressed targets, all of which were downregulated (CCL4, CXCL5, IL12B, IL13, IL17A, IL20, IL5, KITLG, TSLP). However, several of these (IL13, IL20, IL5, TSLP) also showed ≤ 50% detectability in Olink (Figure 1C). This might be attributed to a potential batch effect, as 7 of 8 control samples were run separately from the COVID-19 samples, and the control sample run had higher LODs for most targets versus the COVID-19 sample run (Supplemental Figure 2). The Luminex panel did not uniquely identify any targets. However, when non-specific binding correction was not applied, Luminex uniquely identified IL7 and TNF (Supplemental Figure 1C).

**Figure 3.**
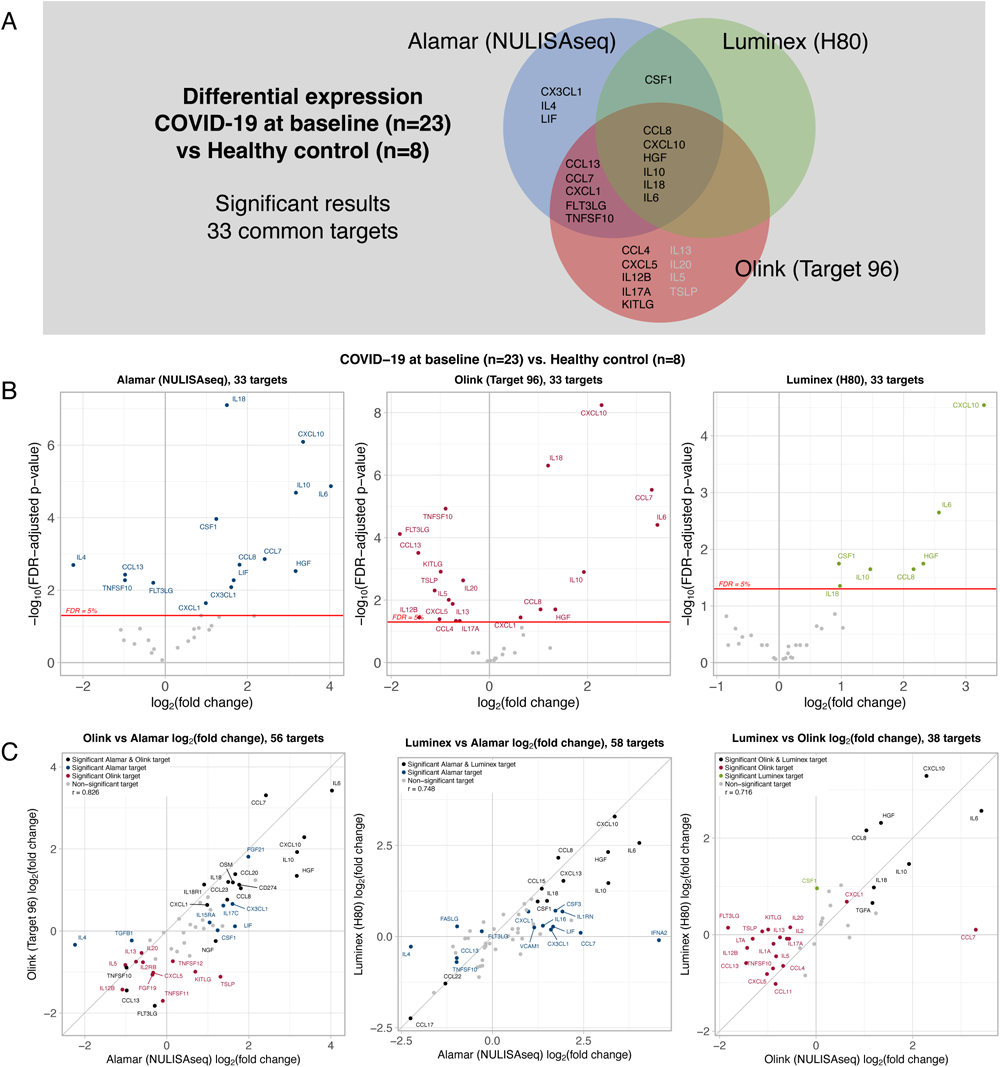
Differential expression comparing inflammatory protein target expression levels for baseline (Visit 1) COVID-19 patients (n=23) and Healthy Controls (n=8). (A) Venn diagram shows overlaps of significant results across panels. Targets in grey text for Olink indicate detectability less than 50%. (B) Volcano plots show -log10 FDR-adjusted p-values versus log2 fold changes for the 33 common targets. (C) Scatterplots show the relationship of log2 fold changes for all common targets in each platform pair.

### Differential expression in COVID-19 patients versus healthy controls, pairwise comparisons for common targets

Alamar NULISAseq and Olink Target 96: Of 56 common targets, Alamar and Olink identified 17 common significant targets (Figure 3C, left). Alamar and Olink log2(fold change) estimates for the 56 targets were positively correlated (r = 0.83). Alamar identified an additional 6 upregulated (CSF1, CX3CL1, FGF21, IL15RA, IL17C, LIF) and 2 downregulated (IL4, TGFB1) targets, and Olink identified 11 additional targets, all of which were downregulated (CXCL5, FGF19, IL12B, IL13, IL20, IL2RB, IL5, KITLG, TNFSF11, TNFSF12, TSLP). Again, several of these (IL2RB, IL20, IL13, IL5, TSLP) showed ≤ 50% detectability in Olink, and their significance may be related to a batch effect.

Alamar NULISAseq and Luminex H80: Of 58 common targets, Alamar and Luminex identified 11 common significant targets (Figure 3C, middle). Alamar and Luminex log2(fold change) estimates for the 58 targets were positively correlated (r = 0.75). Alamar identified an additional 9 upregulated (CCL7, CSF3, CX3CL1, CXCL1, IFNA2, IL16, IL1RN, LIF, VCAM1) and 5 downregulated (CCL13, FASLG, FLT3LG, IL4, TNFSF10) targets, and Luminex identified no additional targets.

Olink Target 96 and Luminex H80: Of 38 common targets, Olink and Luminex identified 7 common significant targets (Figure 3C, right). Olink and Luminex log2(fold change) estimates for the 38 targets were positively correlated (r = 0.72). Olink identified an additional 2 upregulated (CCL7, CXCL1) and 16 downregulated (CCL11, CCL13, CCL4, CXCL5, FLT3LG, IL12B, IL13, IL17A, IL1A, IL2, IL20, IL5, KITLG, LTA, TNFSF10, TSLP) targets, and Luminex identified 1 additional upregulated target (CSF1). Some significant Olink targets showed ≤ 50% detectability (IL13, IL1A, IL2, IL20, IL5, TSLP).

In summary, in pairwise across-platform comparisons of differential expression in COVID-19 at enrollment versus healthy control, Pearson correlation of log2(fold change) estimates were reasonably high and fairly similar, ranging from the highest for Alamar-Olink (r = 0.83) to lowest for Olink-Luminex (r = 0.72). All platform pairs had some findings in common, and each platform showed some unique findings, with Luminex showing the fewest unique findings. In some cases, unique findings corresponded with superior detectability, e.g., Alamar IL4 and LIF (which had ≤ 50% detectability for Olink), and Alamar FASLG and IFNA2 (which had ≤ 50% detectability for Luminex)).

### Differential expression in COVID-19 patients versus healthy controls, full panel results

The Alamar panel detected 84 (41.4%) of 203 targets that were differentially expressed (73 upregulated, 11 downregulated). The Olink panel detected 41 (44.6%) of 92 targets that were differentially expressed (17 upregulated, 24 downregulated). The Luminex panel detected 12 (15%) of 80 targets that were differentially expressed (10 upregulated, 2 downregulated). Thus the Alamar and Olink panels yielded similar rates of differentially expressed targets, while Luminex showed the smallest, and Olink had a larger percentage of downregulated targets. Supplemental Figure 3A shows volcano plots of full-panel differential expression results for each platform.

### Differential expression: Change over time in COVID-19, 33 common targets in 3 assays

For the 33 common targets, 8 showed significant changes over time in COVID-19 patients in all 3 panels (Figure 4A and 4B). Luminex uniquely identified 2 additional targets (IL7, VEGFA); Alamar identified 5 additional targets (CSF1, IL13, IL20, IL4, TNF), and Olink identified 2 (CCL3, KITLG). The non-specific binding correction resulted in 2 more significant targets for Luminex (CCL13, TNFSF10; Supplemental Figure 1C). Figure 5 shows expression over time for selected targets. Supplemental Figure 4 shows all remaining targets that were significant in one or more platforms.

**Figure 4.**
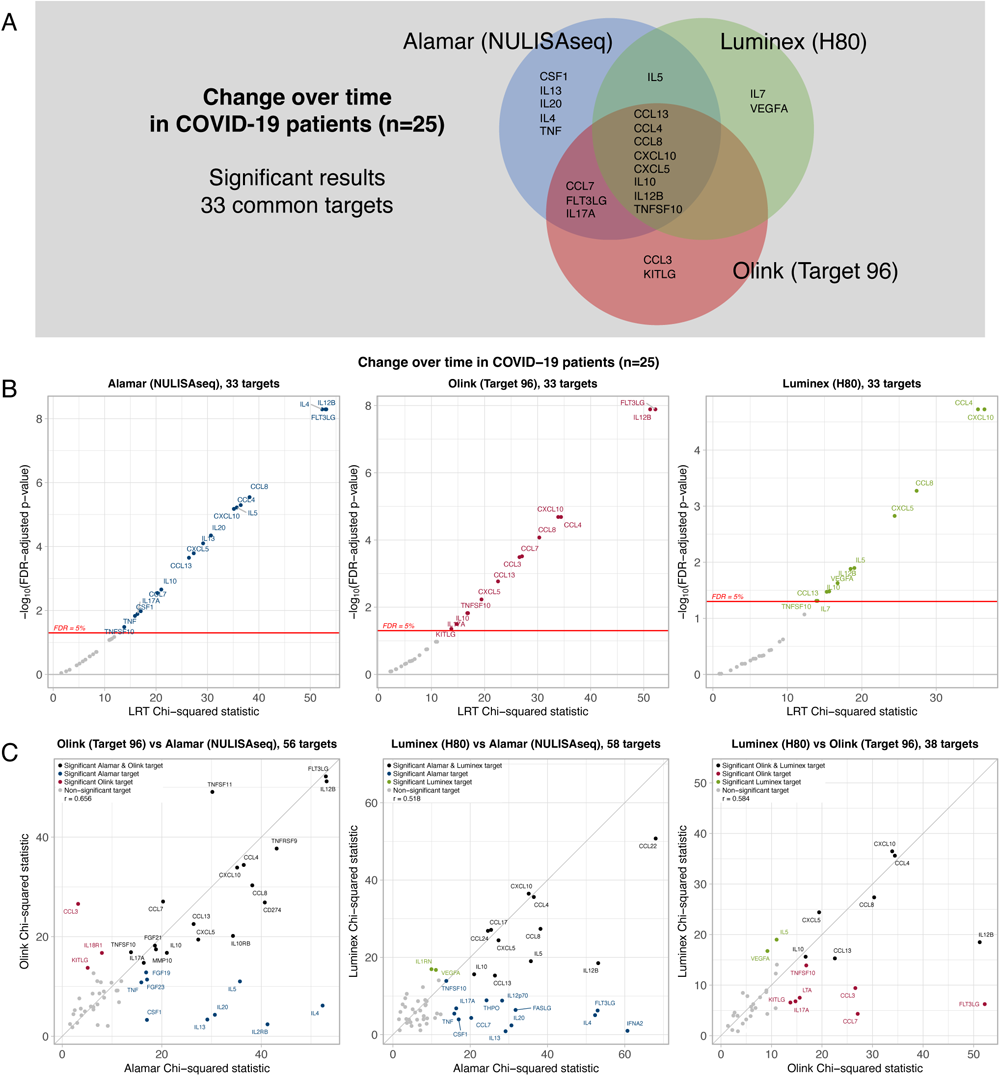
Longitudinal differential expression in COVID patients (n=23). Change over time for inflammatory protein target expression was assessed for the COVID-19 samples using linear mixed effects models with target expression level as outcome, a categorical time fixed effect for each visit, and subject random intercept. The overall significance of the time fixed effect was tested using a likelihood ratio test (LRT). False-discovery rate (FDR) correction was applied to the 33 targets common to all three platforms, and to each subset of overlapping targets for each platform pair. (A) Venn diagram shows overlaps of significant results across platforms. (B) Volcano plots show -log10 FDR-adjusted p-values versus the LRT chi-squared statistic for the 33 common targets. (C) Scatterplots show the relationship of LRT chi-squared statistics for all common targets in each platform pair.

**Figure 5.**
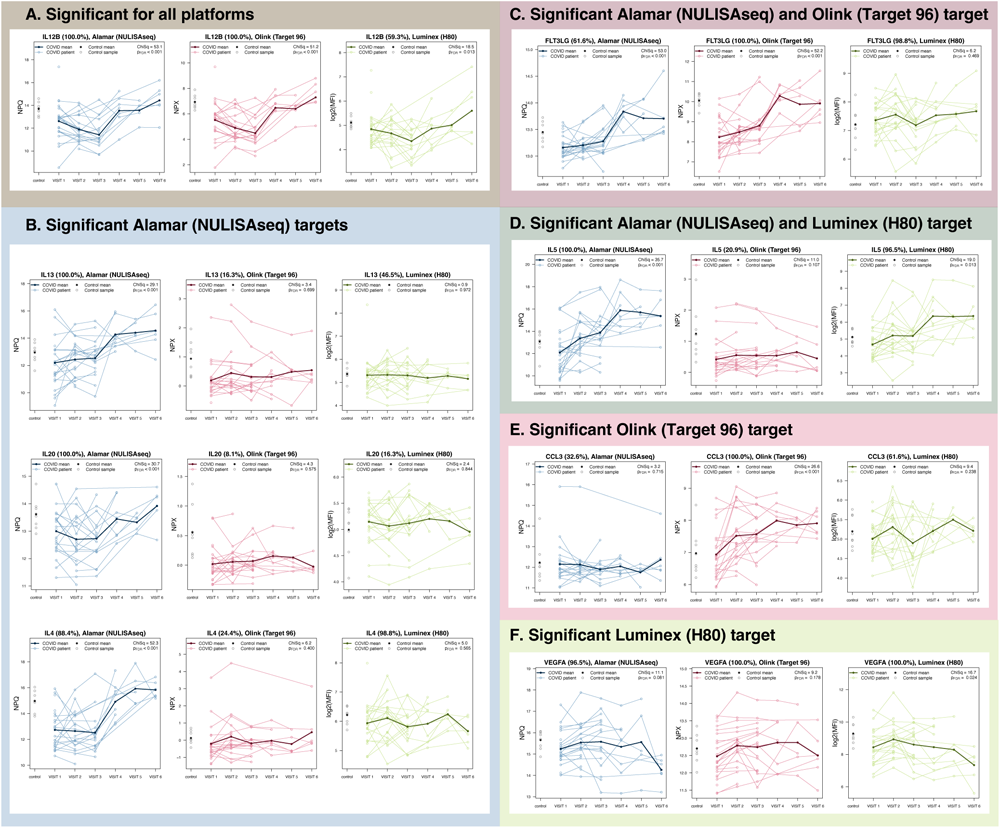
Selected targets showing significant change over time in COVID-19 for one or more platforms. Longitudinal differential expression of inflammatory markers was assessed for 78 serum samples collected from 23 COVID-19 patients over 6 time points. The 6 visits correspond to time of study enrollment (VISIT 1), and days 4, 7, 14, 21, and 28 (VISIT 2-6). Control sample values are shown at the far left side of plots. Values shown in parentheses in plot titles represent target detectability. (A) IL12B was among the targets showing most significant changes over time across all three platforms. (B) Alamar alone showed significant changes in IL13, IL20, and IL4, which are all relatively low-abundance targets that had greater detectability for the Alamar assay, with the exception of IL4 for Luminex. (C) Alamar and Olink both showed significant changes in FLT3LG, but Luminex did not, even though detectability was higher for Luminex than Alamar. (D) Alamar and Luminex both showed significant changes in IL5, but Olink did not. Olink showed low detectability for IL5. (E) CCL3 was only significant for Olink, which also had higher detectability than the other two platforms. (F) VEGFA was only significant for Luminex, despite high detectability for all three platforms.

### Differential expression: Change over time in COVID-19, pairwise comparisons for common targets

Alamar NULISAseq and Olink Target 96: Of 56 shared targets, Alamar and Olink identified 17 common significant targets (Figure 4C, left). Alamar and Olink chi-squared statistics for the 56 targets were positively correlated (r = 0.66). Alamar identified 9 additional targets (CSF1, FGF19, FGF23, IL13, IL20, IL2RB, IL4, IL5, TNF), and Olink identified 3 additional targets (CCL3, IL18R1, KITLG). Notably, IL13, IL20, IL2RB, IL4, and IL5 had ≤ 50% detectability for Olink, and CCL3 had ≤ 50% detectability for Alamar.

Alamar NULISAseq and Luminex H80: Of 58 common targets, Alamar and Luminex identified 11 common significant targets (Figure 4C, middle). Alamar and Luminex chi-squared statistics for the 58 targets were positively correlated (r = 0.52). Alamar identified 13 additional targets (CCL7, CSF1, FASLG, FLT3LG, IFNA2, IL12p70, IL13, IL17A, IL20, IL4, THPO, TNF, TNFSF10), and Luminex identified 2 additional targets (IL1RN, VEGFA). Notably, IL20 and FASLG had ≤ 50% detectability for Luminex versus 100% for Alamar; IL17A had 53.4% detectability for Luminex versus 94.2% for Alamar; IL12p70 had 58.1% detectability for Luminex versus 87.2% for Alamar.

Olink Target 96 and Luminex H80: Of 38 common targets, Olink and Luminex identified 7 common significant targets (Figure 4C, right). Olink and Luminex chi-squared statistics for the 38 targets were positively correlated (r = 0.58). Olink identified 7 additional targets (CCL3, CCL7, FLT3LG, IL17A, KITLG, LTA, TNFSF10), and Luminex identified 2 additional targets (IL5, VEGFA). Notably, IL17A had higher detectability for Olink (83.7%) than Luminex (53.5%), and IL5 had higher detectability for Luminex (97.7%) than Olink (20.9%).

In summary, in pairwise across-platform comparisons of differential expression over time in COVID-19 patients, correlation of LRT chi-squared statistics was highest for Alamar-Olink (r=0.66), and these two platforms had the largest number of common significant findings (17 targets). Luminex identified the least number of significant targets in the pairwise comparisons, and several targets that Luminex missed also had low detectability for Luminex. In each comparison, Alamar identified more significant changes over time than the other platform.

### Differential expression: Change over time in COVID-19, full panel results

Considering the full panels for each platform, the Alamar panel detected a total of 86 (42.4%) of 203 targets showing significant change over time, the Olink panel detected a total of 29 (31.5%) of 92 targets showing significant change over time, and the Luminex panel detected a total of 15 (18.8%) of 80 targets showing significant change over time. Supplemental Figure 3B shows volcano plots of full-panel differential expression results for each platform.

### Logistic regression: Predicting COVID-19 outcome using protein expression levels at enrollment

Due to small sample size (9 least severe versus 14 most severe), none of the panels showed significant FDR-adjusted results. Based on unadjusted p < 0.05 significance, Alamar (NULISAseq) showed 1 target with odds ratio significantly below one (CCL3) and 6 targets with odds ratio significantly above one (GDF15, IL12p70, IL17B, IL35, MDK, S100A9) for the most versus least severe trajectory COVID-19 patients. However, CCL3 and IL35 had low detectability for Alamar (32.6% and 5.8%, respectively). For the other targets, area under the curve (AUC) ranged from 0.833 to 0.746. Olink (Target 96) and Luminex (H80) showed no significant targets. However, for Luminex without non-specific binding correction, THPO odds ratio was significantly above 1, and VCAM1 had a large effect size (odds ratio=1470, p=0.06, AUC=0.803). Supplemental Figure 3C shows volcano plots and ROC curves.

Detailed results, including detectability, inter-assay correlation, and differential expression results for each assay and target can be found in Supplemental Data A.

## Discussion

In this study, we conducted the first cross-platform comparison to evaluate the measurement of inflammatory markers in COVID-19 samples using three advanced and highly sensitive technologies: bead-based immunoassay Luminex H80, proximity extension assay (PEA) Olink Target 96, and a novel proximity ligation assay (PLA), Alamar NULISAseq. The primary objectives of this study were twofold: firstly, to provide insights into platform selection for future investigations; and secondly, to enhance the interpretation of previous research findings by establishing a comprehensive understanding of the immunoassays employed for measuring the inflammatory-related proteome in the context of COVID-19.

The Alamar, Olink, and Luminex panels are complementary in that they each include a unique set of markers. This study highlights strengths and weaknesses of the overall panels and for common marker subsets. The observed differences in detectability between platforms illustrate assay-specific variations in sensitivity, emphasising the importance of considering this assay characteristic when evaluating protein abundance. We found that overall detectability for the platforms was highest for Alamar, followed by Olink and then Luminex.

Larger differences in detectability generally corresponded to lower correlation of protein measures across platforms. However, there were some exceptions where detectability was high in both platforms and yet correlation was low. For example, CSF1 showed 100% detectability by both Alamar and Olink, but Kendall coefficient was 0.05. In general, Alamar and Olink correlation tended to be greater than correlation of either platform with Luminex.

In many cases, detectability differences also translated to differences in differential expression findings. For example, Alamar showed the highest detectability for IL13 and IL20, and was the only platform able to detect significant increases in these targets over time in COVID-19 patients. Alamar and Luminex both showed higher detectability than Olink for IL5, and thus were able to show significant increase in IL5 over time. Increasing trends for IL13 and IL5 have been previously reported in severe COVID-19 (17), and IL13 has been implicated as a driver of COVID-19 severity (18). Similarly, the higher detectability of Alamar (92.9%) for IFNA2 than Luminex (40.0%) enabled Alamar to demonstrate statistically significant changes for IFNA2 in baseline and longitudinal comparisons, as expected for an antiviral response (19).

However, high detectability did not always correspond to significant findings. For example, Alamar alone was able to detect differences at baseline and increase over time in IL4, which had low detectability in Olink (24.4%), but 97.7% detectability in Luminex. IL4 increases have also been noted previously in severe COVID-19 (17). Furthermore, Luminex did not show changes in FLT3LG even with detectability 98.8%, while both Alamar (60.9% detectability) and Olink (100% detectability) showed significant FLT3LG baseline differences and changes over time in COVID-19. This shows that high detectability does not guarantee the ability to detect meaningful biological differences.

Low inter-platform correlation for targets with high detectability and incongruent differential expression findings for these targets might result from differences in dynamic range between platforms. Even when samples are above LOD, a smaller dynamic range will limit the ability to detect true biological variation. Additionally, incongruent findings could be due to differences in specificity among platforms. While analytical sensitivity relates to the LOD, analytical specificity depends on discrimination between targets and other potentially interfering proteins (20). Non-specific signal could potentially obscure biological differences.

The assay platforms studied here rely on some shared mechanisms for ensuring specificity, such as dual antibody recognition. However, they also employ various different strategies to achieve specificity. For Luminex we applied a non-specific binding correction utilising Assay Chex beads (21). Overall Luminex detectability was enhanced through the application of this method (Supplemental Figure 1), as the method estimated and removed non-specific signal on a per-well basis, including the negative control wells used to define LOD. Differential expression analysis after the correction resulted in more findings for some comparisons and fewer for others. Conversely, in the NULISA, the implementation of capture and release steps was designed to yield a significant reduction of non-specific binding.

Our study reinforces previous findings in terms of cytokines associated with COVID-19. For example, all 3 platforms showed upregulated IL6, CXCL10, and IL10 at baseline, and these have been identified as an important triad of plasma markers that distinguish more severe COVID-19 patients (22). Furthermore, CCL8, CXCL10, IL18, and IL6 (identified by all 3 platforms), and CCL7 (identified by Alamar and Olink) were found to be consistently upregulated in COVID-19 versus controls in a meta-analysis of five clinical studies (23). Many of the targets identified by the Alamar panel to be predictive of COVID-19 outcomes have been previously associated with outcome severity in COVID-19, e.g., GDF15 (24, 25), S100A9 (26), IL17B (27), MDK (28), and IL12p70 (29, 30).

Cross-platform comparisons showed that Olink identified the most differentially expressed targets at baseline, although for low detectability targets this was likely confounded by a batch effect, followed by Alamar and then Luminex. In longitudinal analyses, Alamar identified the most targets, followed by Olink and then Luminex.

While this study reveals similarities and differences for the 3 assays, it is worth noting some limitations. Despite normalization methods applied to reduce bias, data suggests that some batch effect is still present for the Olink data. Specifically, Olink found more significantly downregulated targets in the baseline COVID-19 versus healthy control comparison relative to the 2 other platforms, and several of these targets showed low detectability. Observed differences in background between the two Olink runs could potentially explain this. In standard practice, it makes sense to exclude such low detectability targets from further analyses, but for the sake of inter-platform comparisons we opted to include them. Also note that while Olink found baseline differences for IL5, IL13, or IL20, Olink did not show longitudinal findings for these while Alamar did, as did Luminex for IL5. In spite of this, many Olink baseline differential expression significant findings were still in agreement with other platforms. The longitudinal analysis for Olink is not affected by any batch effects, as all COVID-19 samples were run together.

In addition to performance-related criteria such as sensitivity and specificity, other key factors to consider when selecting an immunoassay platform include panel coverage and ease of use. Panel coverage emerges as a pivotal criterion for platform selection. The Alamar panel encompasses over 200 targets, making it a good choice for researchers seeking a wide range of analytes. Olink, while providing 92 targets in its standard offering, presents the potential for comprehensive exploration through the Olink Explore option, which includes more than 3000 targets. Luminex, although starting with fewer targets, stands out for its flexibility and customization capabilities, allowing researchers to run partial plates and to tailor panels according to specific research needs. Thus, the choice of platform can be aligned with the desired scope and specific targets of interest. Ease of use also plays a crucial role in the selection process. The Alamar platform excels in user-friendliness, offering a fully integrated assay process that spans from setup (achievable in under 30 minutes) to the pooling of NGS libraries (completed in less than 6 hours, albeit NGS runs separately). The input volume requirement of 10 µL enhances its accessibility. Olink and Luminex are both more manual and require similar hands-on time to process samples. While Olink has an input volume of only 1 µl, Luminex requires the highest volume at 25 µl. Researchers should weigh the convenience of assay setup and run times based on their specific project requirements.

In this study, we aimed to comprehensively assess the detectability, correlation, and differential expression of protein targets using three different assays: Luminex, Olink, and Alamar. Despite the noted limitations, our findings illuminate aspects of protein quantification within the context of COVID-19, and provide insights into the commonalities and distinctions among these platforms. Specifically, our results reveal that overall detectability for the platforms was highest for the Alamar assay, followed by Olink and Luminex. Additionally, Alamar and Olink showed the greatest concordance in pairwise platform comparisons. Although most of the differences in cross-platform correlation and differential expression results could be explained by differences in sensitivity of different platforms, some might also be attributable to differences in assay dynamic range or specificity. This study thus shows how the diverse characteristics of these platforms have potential implications for the accuracy and robustness of downstream analyses. Only with the combination of high sensitivity, high specificity, and a sufficiently broad dynamic range can biological insights be reliably obtained.

## Acknowledgements

We thank the IMPACC network and IMPACC patients for access to samples.

## Footnotes

1 Funded by grant 2U19AI057229 from the National Institutes of Health

