## Supplemental Figures for "Cross-platform comparison of highly-sensitive immunoassays for inflammatory markers in a COVID-19 cohort^1^"

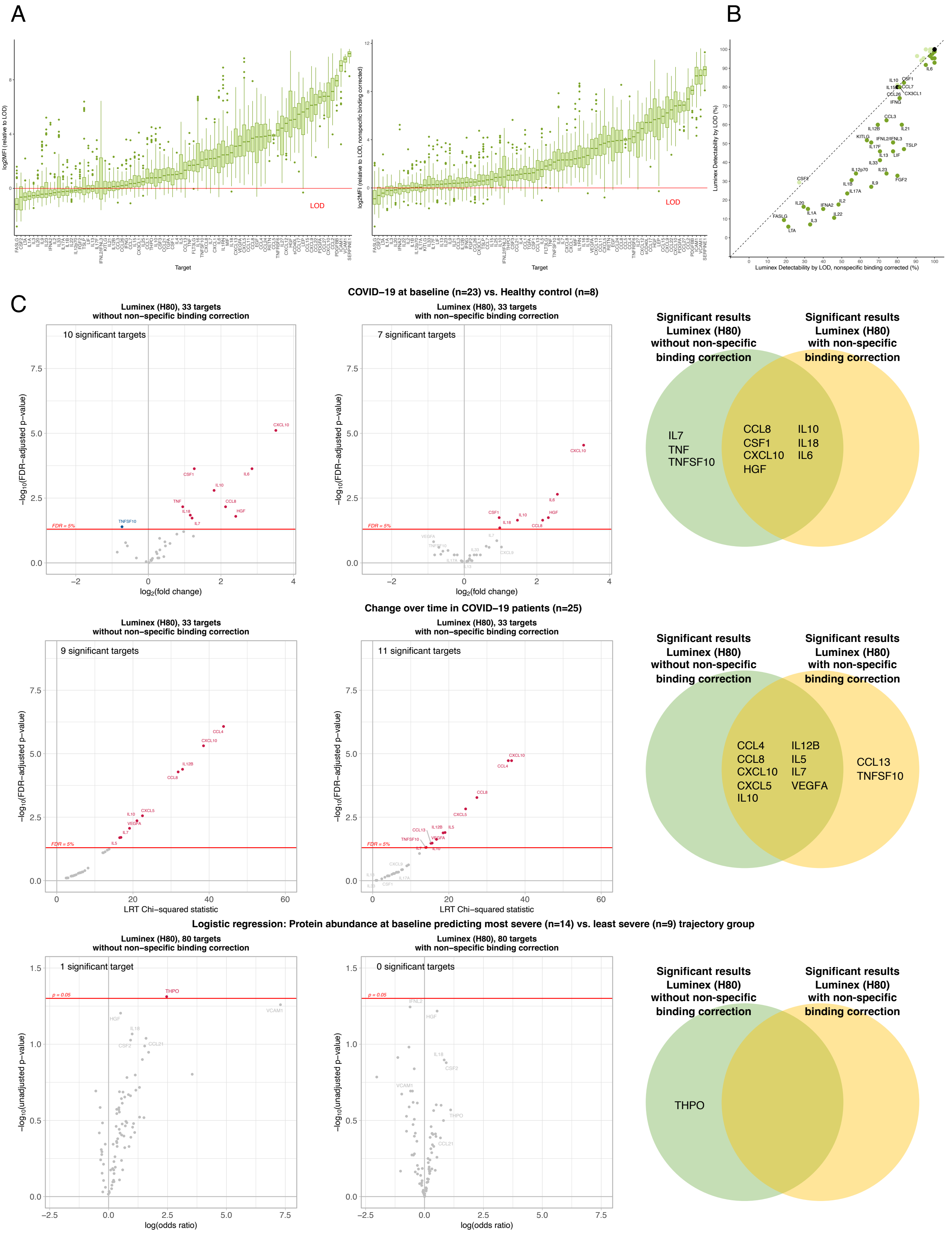
A
B
C
log2MFI (relative to LOD)
log2MFI (relative to LOD, nonspecific binding corrected)
LOD
LOD
Target
Target
Luminex Detectability by LOD (%)
Luminex Detectability by LOD, nonspecific binding corrected (%)
COVID−19 at baseline (n=23) vs. Healthy control (n=8)
Luminex (H80), 33 targets
without non−specific binding correction
Luminex (H80), 33 targets
with non−specific binding correction
10 significant targets
7 significant targets
−log10(FDR−adjusted p−value)
log2(fold change)
FDR = 5%
CXCL10
CSF1
IL10
IL6
TNF
IL18
CCL8
HGF
IL7
TNFSF10
Significant results
Luminex (H80)
without non-specific
binding correction
Significant results
Luminex (H80)
with non-specific
binding correction
IL7
TNF
TNFSF10
CCL8
CSF1
CXCL10
HGF
IL10
IL18
IL6
Change over time in COVID−19 patients (n=25)
9 significant targets
11 significant targets
LRT Chi−squared statistic
CCL4
CXCL10
IL12B
CCL8
CXCL5
VEGFA
IL5
CCL13
TNFSF10
CCL4
CCL8
CXCL10
CXCL5
IL10
IL12B
IL5
IL7
VEGFA
CCL13
TNFSF10
Logistic regression: Protein abundance at baseline predicting most severe (n=14) vs. least severe (n=9) trajectory group
Luminex (H80), 80 targets
without non−specific binding correction
Luminex (H80), 80 targets
with non−specific binding correction
1 significant target
0 significant targets
−log10(unadjusted p−value)
log(odds ratio)
p = 0.05
THPO
VCAM1
HGF
CSF2
CCL21
THPO

**Supplemental Figure 1.** Non-specific binding correction in Luminex H80 assay. (A) Signal relative to LOD with (right) and without (left) non-specific binding correction. (B) Comparison of detectability before (y-axis) and after (x-axis) non-specific binding correction. Among the targets subjected to non-specific binding correction, 11 exhibited lower detectability (light green), 31 showed equal detectability (black), and 38 displayed higher detectability (dark green) compared to the detectability without non-specific binding correction. (C) Differential expression results with and without non-specific binding correction in Luminex H80 assay.

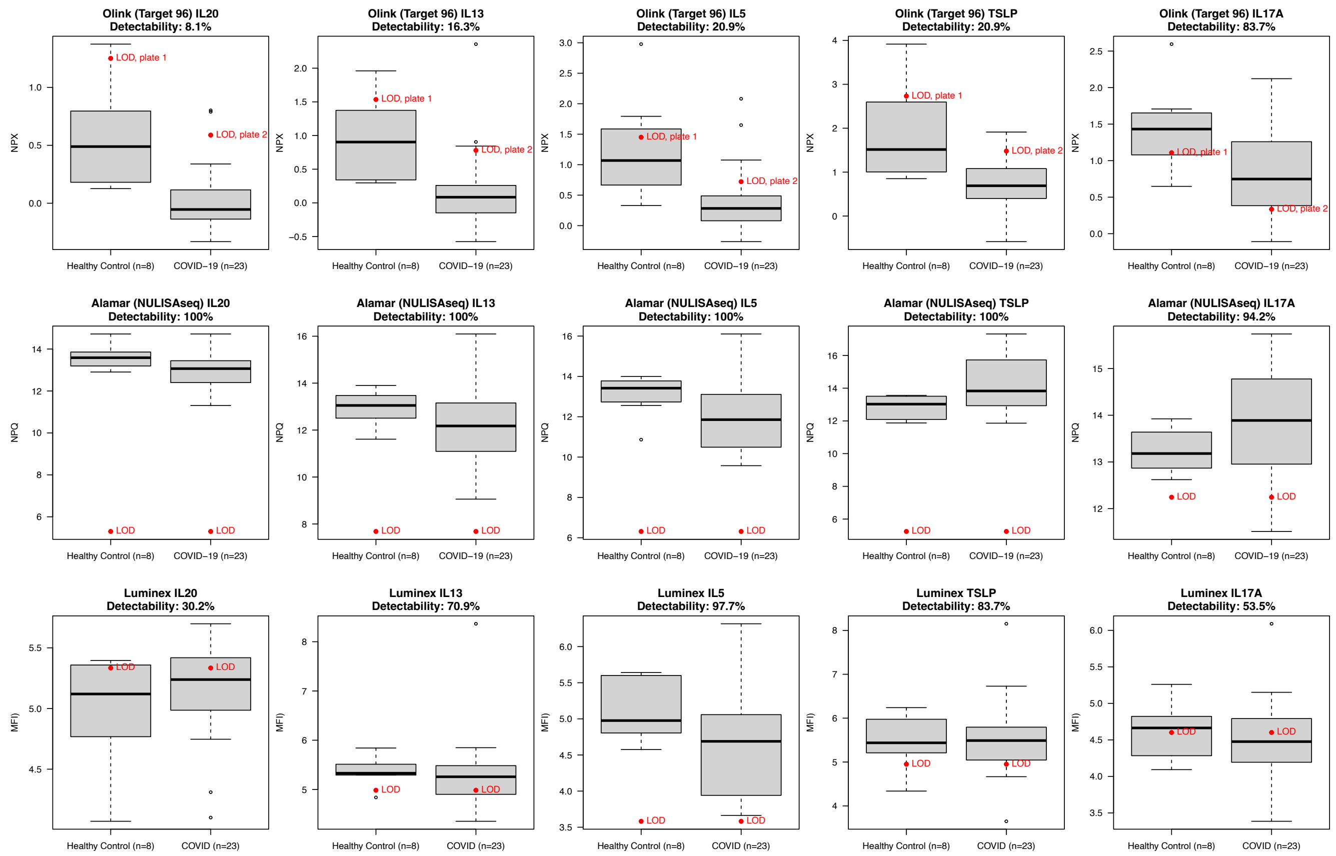


**Supplemental Figure 2.** In the comparison of baseline protein abundance levels for (Visit 1) COVID-19 patients (n=23) versus Healthy Controls (n=8) across the 33 protein targets common to all 3 platforms, Olink (Target 96) assay detected more downregulated differentially expressed targets for COVID-19 than the other platforms. However, 5 of the unique downregulated Olink targets had below 100% detectability (shown in boxplots in top row, above), and 4 of these had below 25% detectability. This could be driven by a potential run-related batch effect in the Olink data. Since 7 of 8 control samples were run on a separate plate, and the 2 runs had different background levels (i.e., limit of detection; LOD) after normalization, the differential expression for these downregulated, low-detectability targets in the Olink assay may be driven by the difference in background levels. Target distributions for Alamar and Luminex assays, are shown in the middle and bottom rows, respectively. For Alamar and Luminex assays, COVID-19 and control samples were run together on the same plate, so LODs are the same across groups and there are no potential run-related batch effects.

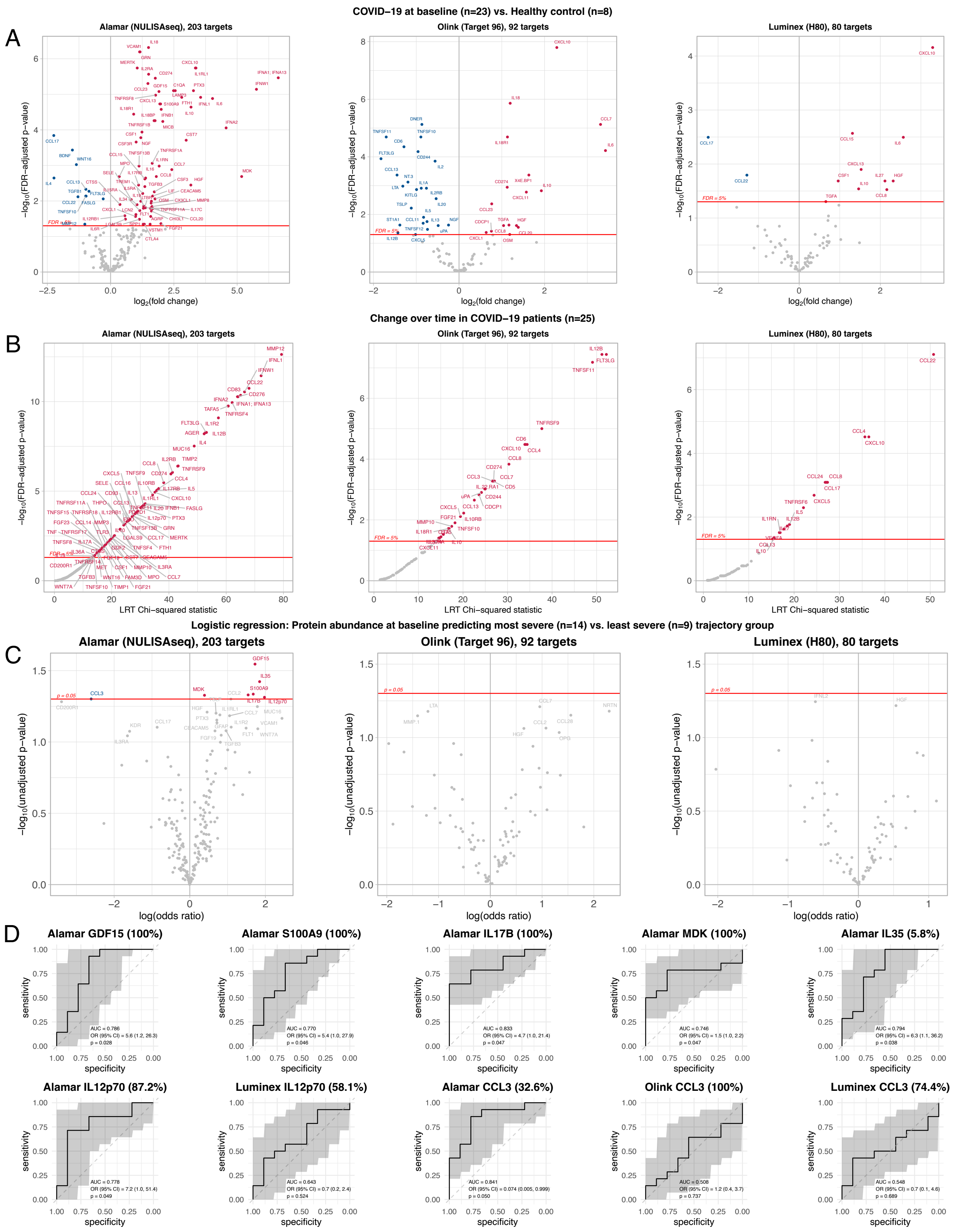
COVID−19 at baseline (n=23) vs. Healthy control (n=8)
A
Alamar (NULISAseq), 203 targets
Olink (Target 96), 92 targets
Luminex (H80), 80 targets
−log10(FDR−adjusted p−value)
log2(fold change)
FDR = 5%
Change over time in COVID−19 patients (n=25)
B
LRT Chi−squared statistic
Logistic regression: Protein abundance at baseline predicting most severe (n=14) vs. least severe (n=9) trajectory group
C
−log10(unadjusted p−value)
log(odds ratio)
p = 0.05
D
Alamar GDF15 (100%)
Alamar S100A9 (100%)
Alamar IL17B (100%)
Alamar MDK (100%)
Alamar IL35 (5.8%)
Alamar IL12p70 (87.2%)
Luminex IL12p70 (58.1%)
Alamar CCL3 (32.6%)
Olink CCL3 (100%)
Luminex CCL3 (74.4%)
sensitivity
specificity

**Supplemental Figure 3.** (A) Differential expression comparing protein target expression levels for baseline (Visit 1) COVID-19 patients (n=23) and Healthy Controls (n=8). Results are shown for all targets in each platform. The Alamar panel detected 73 upregulated and 11 downregulated targets, for an overall total of 84 (41.4%) of 203 targets differentially expressed. The Olink panel detected 17 upregulated and 24 downregulated targets, for an overall total of 41 (44.6%) of 92 targets differentially expressed. The Luminex panel detected 10 upregulated and 2 downregulated targets, for an overall total of 12 (15%) of 80 targets differentially expressed. (B) Differential expression: COVID-19 patients change over time. Showing results for all targets for each platform. The Alamar panel detected a total of 86 (42.4%) of 203 targets showing significant change over time. The Olink panel detected a total of 29 (31.5%) of 92 targets showing significant change over time. The Luminex panel detected a total of 15 (18.8%) of 80 targets showing significant change over time. (C) Logistic regression: Predicting trajectory group severity outcome based on protein expression at enrollment. An exploratory analysis was carried out to assess whether protein expression levels at baseline could predict the most severe (trajectory groups 4 and 5; n=14) versus the least severe (trajectory groups 1, 2, and 3; n = 9) outcomes for COVID-19 patients. Results are shown for all targets for each platform. No platforms showed any FDR-corrected results, so unadjusted p-values are shown. Significance was set at unadjusted $p < 0.05$. Targets with $p < 0.1$ are labelled. Alamar (NULISAseq) was the only panel that showed significant targets. (D) Receiver operating characteristic (ROC) curves for significant targets. ROC curve plots were generated using R package pROC (Xavier Robin, Natacha Turck, Alexandre Hainard, Natalia Tiberti, Frédérique Lisacek, Jean-Charles Sanchez and Markus Müller (2011). pROC: an open-source package for R and S+ to analyze and compare ROC curves. BMC Bioinformatics, 12, p. 77. DOI: 10.1186/1471-2105-12-77 <http://www.biomedcentral.com/1471-2105/12/77/>).

### Change over time in COVID-19 patients

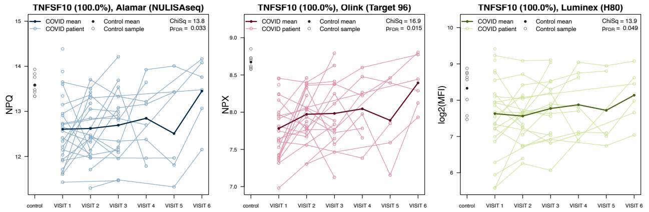


#### Significant for all platforms

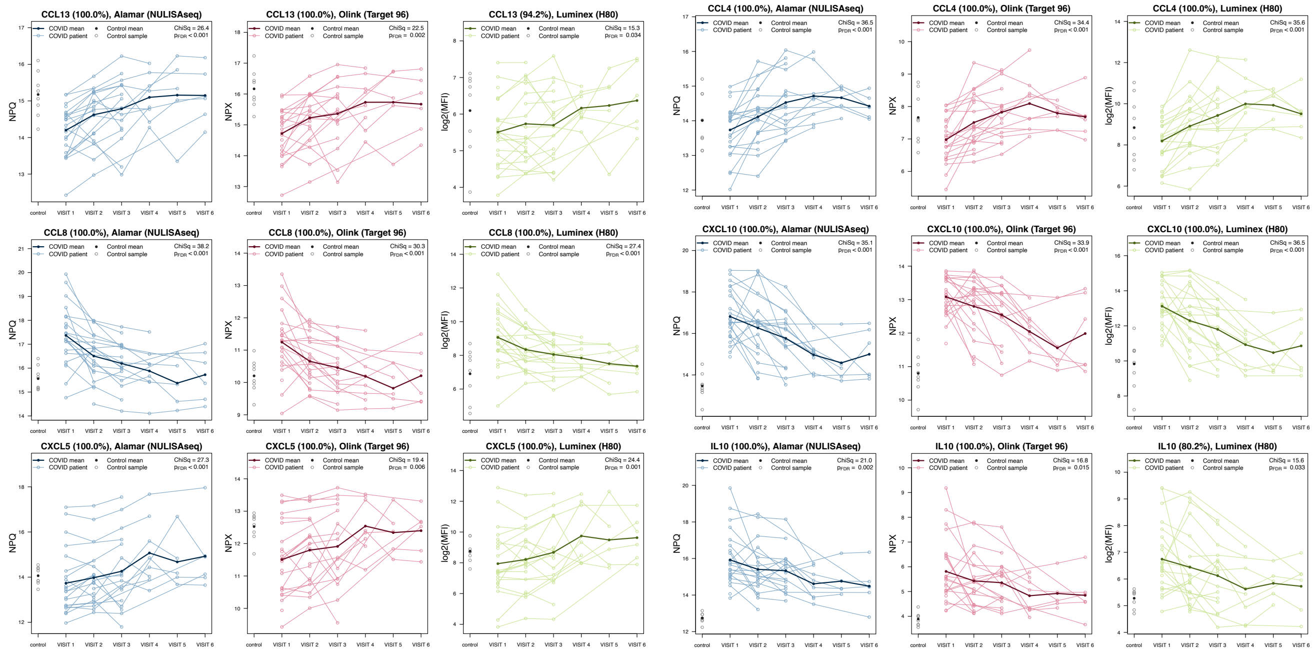


#### Significant for Alamar (NULISAseq) and Olink (Target 96)

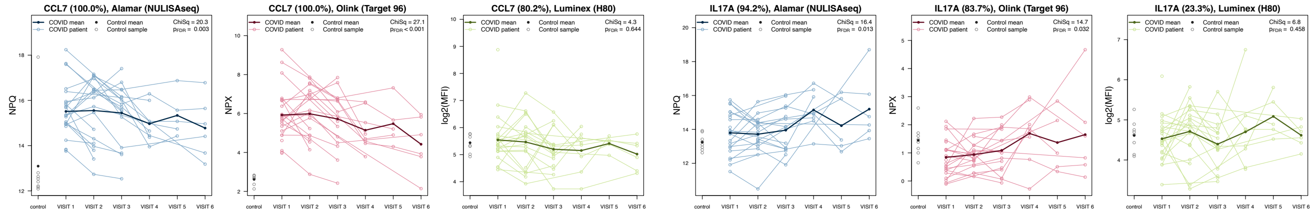


#### Significant for Luminex (H80)

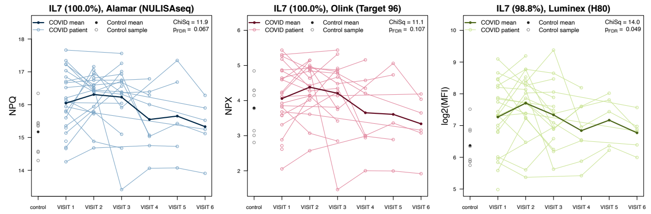


#### Significant for Olink (Target 96)

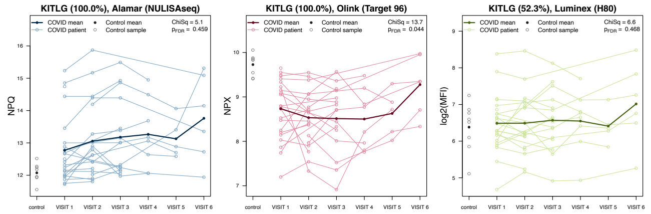


#### Significant for Alamar (NULISAseq)

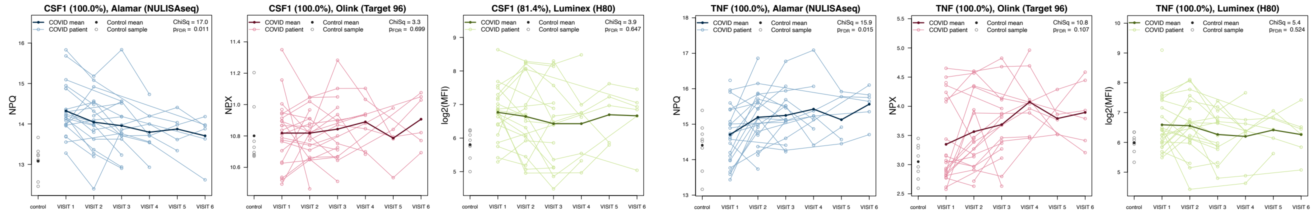


40

**Supplemental Figure 4.** Trajectory plots for additional targets showing significant change over time in COVID-19 patients for one or more platforms. Control sample values are shown at the far left side of plots. The 6 visits correspond to time of study enrollment (VISIT 1), and days 4, 7, 14, 21, and 28 (VISIT 2-6). Values shown in parentheses in plot titles represent target detectability.
